# Calculating the optimal hematocrit under the constraint of constant cardiac power

**DOI:** 10.1101/2020.06.09.141374

**Authors:** M. Sitina, H. Stark, S. Schuster

## Abstract

In humans and higher animals, a trade-off between sufficiently high erythrocyte concentrations to bind oxygen and sufficiently low blood viscosity to allow rapid blood flow has been achieved during evolution. Optimal hematocrit theory has been successful in predicting hematocrit values of about 0.3 - 0.5, in very good agreement with the normal values observed for humans and many animal species. However, according to those calculations, the optimal value should be independent of the mechanical load of the body. This is in contradiction to the exertional increase in hematocrit observed in some animals called natural blood dopers and to the illegal practice of blood boosting in high-performance sports. In this study, we calculate the optimal hematocrit under two different constraints - under a constant driving pressure and under constant cardiac power – and show that the optimal hematocrit under constant cardiac power is higher than the normal value, ranging from 0.5 to 0.7. We use this result to explain the tendency to better exertional performance at an increased hematocrit.

**Statement of Significance:** In humans and higher animals, erythrocytes comprise a volume fraction (hematocrit) of 30-50 % of the blood. Mathematical calculations based on the assumption of constant blood pressure show that the optimal hematocrit value is indeed in that range. However, this optimum should apply to both rest and physical exertion, which is in contradiction to the increase in hematocrit observed in some animals called natural blood dopers and to the illegal practice of blood boosting in sports. Here, we calculate the optimal value based on the alternative constraint of constant cardiac power. We show that this results in a higher optimal value, ranging from 50 to 70 %. In this way, we explain the better exertional performance at an increased hematocrit.

## 1. Introduction

In the blood flow of humans and higher animals, a trade-off between sufficiently high erythrocyte concentration (known as hematocrit) to transport oxygen and sufficiently low blood viscosity to allow rapid flow has been achieved during evolution. Based on the trade-off between these two factors, authors from our group calculated the optimal hematocrit value maximizing oxygen supply to be 0.3-0.5, which is in very good agreement with values observed in humans and many animal species (1, 2). The exact predicted value depends on the formula used to express viscosity in terms of hematocrit. Several other detailed theoretical modelling studies (3, 4) yield the optimal hematocrit value of about 0.4 as well. This shows that even a highly simplified model (1, 2) can lead to relevant results. An optimal resting hematocrit of about 0.4 was also found in an experiment with dogs in which the hematocrit was varied by blood exchange (5).

However, according to the calculations (1-4), the optimal value should be independent of the level of exertion. This is in contradiction to the optimal hematocrit value of 0.5-0.6 found during a blood perfusion of isolated working muscle at a constant perfusion pressure (6). A lower sensitivity of blood flow to hematocrit in working muscle in comparison to resting muscle was also found in (7). Another contradiction comes from the observation in endurance runner animals like dogs or horses, which are called natural blood dopers (8) because they increase their hematocrit at exertion via expulsion of concentrated blood from the spleen. Finally, a counter-example is also the prohibited practice of blood boosting in high-performance sports. With the use of blood boosting, several outstanding records were set and many studies have demonstrated its positive performance effect (9-21). In other studies (22), however, no increase was observed in performance. Even placebo effects have been mentioned to solve this “hematocrit paradox” and the topic remains controversial (8). A drawback of the idea about a positive effect of increased hematocrit is the absence of a clear theoretical explanation: why should the increased hematocrit promote performance, when the theoretical models (1, 3, 4) predict the optimum to be at normal values even at exertion.

Experimental studies using erythropoietin indicate that an increase in hematocrit leads to a weak increase both in the pressure difference and in cardiac power (8, Table 1). In this study, we perform a calculation of the optimal hematocrit under two different, simplifying constraints - under constant perfusion pressure and under constant cardiac power, to check which of these describes observations better. Blood pressure is a well-regulated quantity at rest. At exertion, however, the body switches to performance and takes the risk of increasing blood pressure. The constraint of constant cardiac power implies that at lower blood flow velocities (due to higher blood viscosity), the driving force of the heart increases, so that the product of both remains constant. We show that the resulting optimal hematocrit is higher than the normal value. With this result, we explain the tendency to better exertional physical performance at an increased hematocrit, at least over shorter periods.

**TABLE 1.**
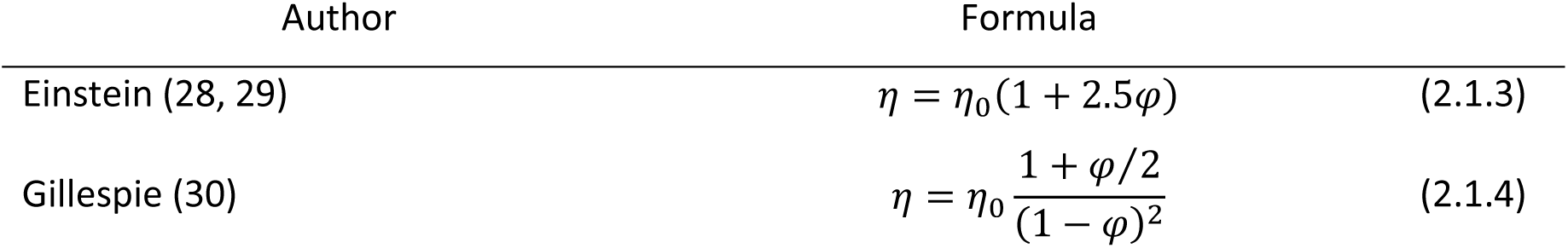
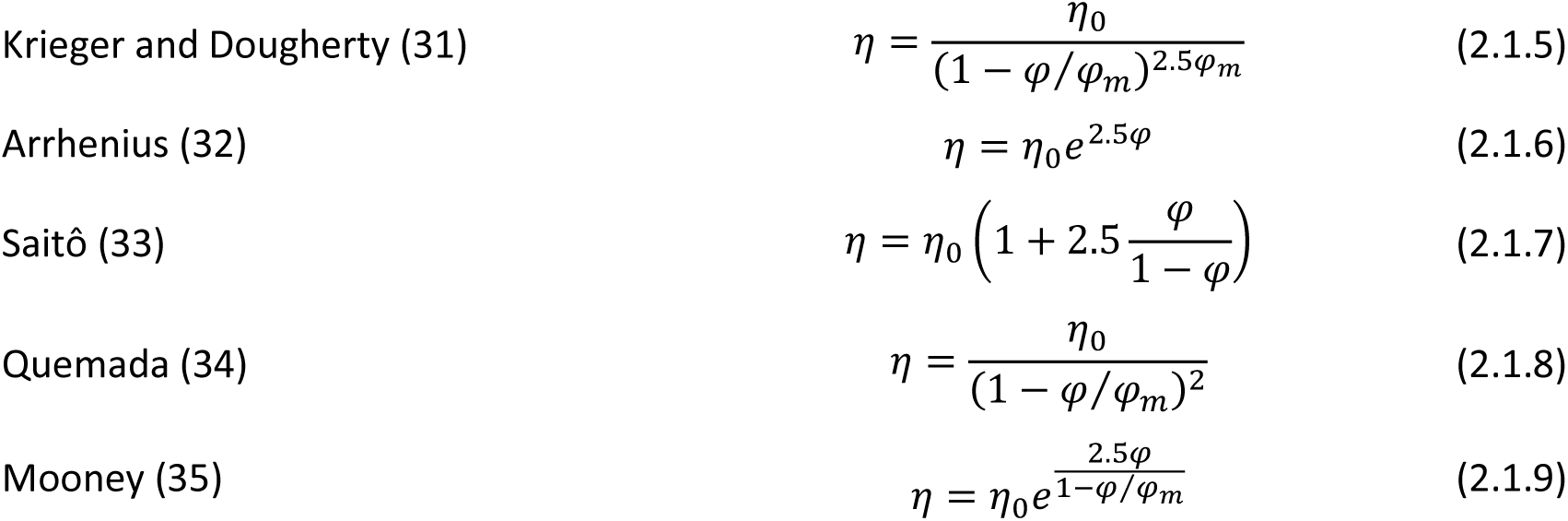
Relationships between blood viscosity *η* and the hematocrit *φ*. *η*_0_ is the viscosity of blood plasma, *φ*_m_ is the maximum possible volume fraction of erythrocytes.

## 2. Methods and Results

### 2.1. Basic equations

According to the Hagen-Poiseuille law (23) the flow *Q* through a tube of length *l* and radius *r* under the pressure difference Δ*p* (alternatively called perfusion pressure or driving pressure) is given by

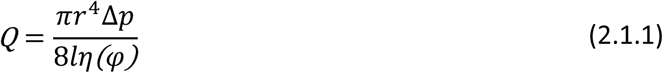

where *η*(*φ*) denotes the fluid viscosity, which is, in the case of blood, a function of the hematocrit *φ*. For our calculations, it is only relevant that the flow is proportional to Δ*p* and inversely proportional to the viscosity *η*. This also holds for an ideal flow in a vessel with elliptical or rectangular cross-section (24) or the flow through porous media (25). Thus, Eq. 2.1.1 can be rewritten as

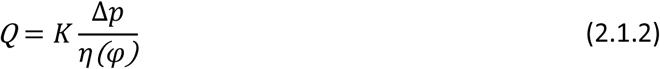

with some constant *K* dependent only on the tube geometry. This relationship can be generalized for the whole circulation or segments of circulation (26, 27), with *Q* being the total flow through the circulation (i.e. cardiac output) and Δ*p* the pressure difference between both ends of the circulation (i.e. the difference between the mean arterial pressure and the pressure in the right atrium of the heart).

In the following sections, we use several relationships between the hematocrit and blood viscosity that are summarized in Table 1. Some of these relationships are illustrated in Figure 1. *η*_0_ denotes the viscosity of the particle-free liquid, which is blood plasma in our case. The hematocrit *φ* represents the volume fraction of erythrocytes in the blood, *φ*_m_ is the maximum possible volume fraction of erythrocytes (maximal packing density). In case of stiff spherical particles, for example, *φ*_m_ equals 74 %. The formulas are discussed in Stark & Schuster (1, 2).

**FIGURE 1.**
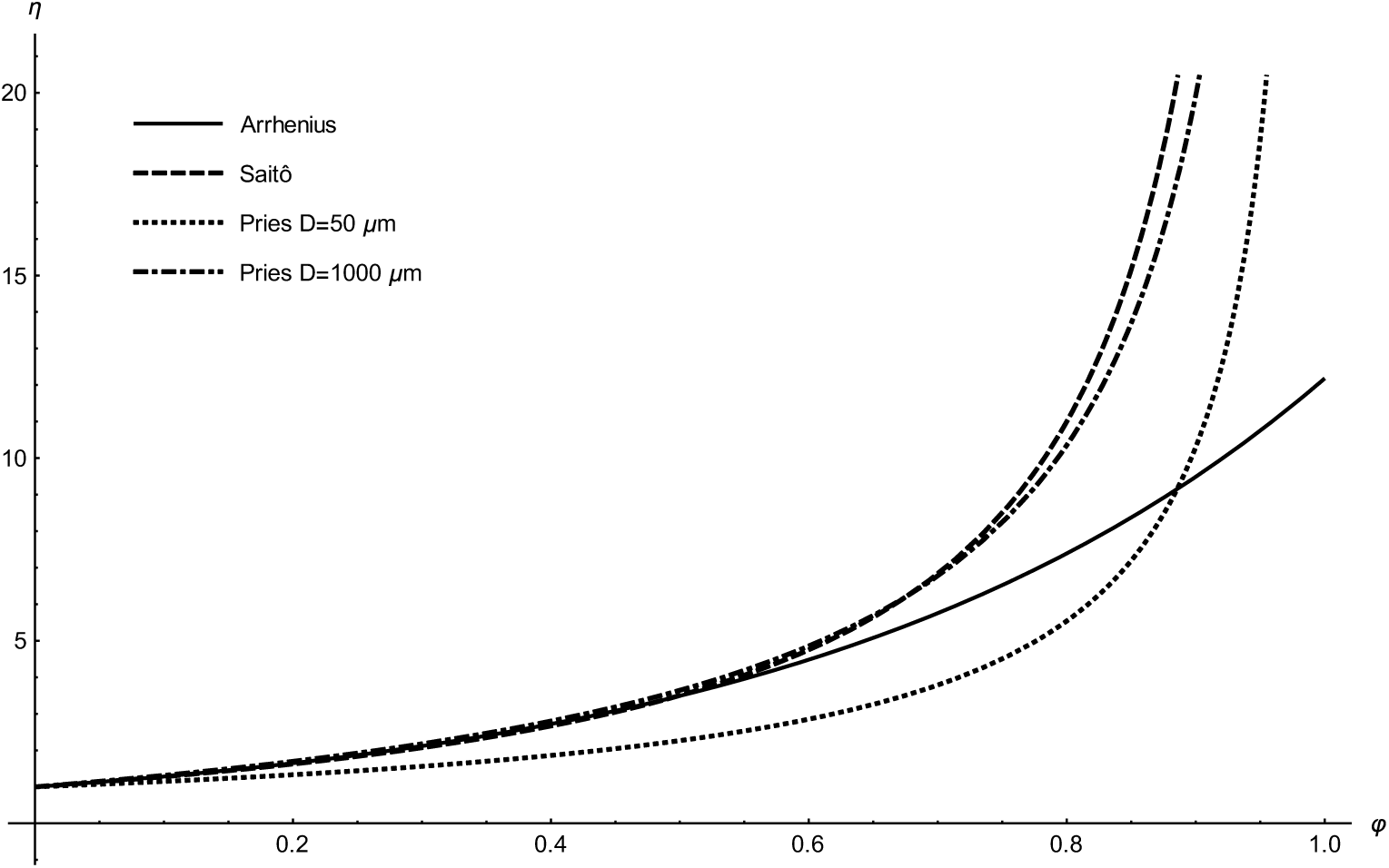
Dependencies of the (relative) viscosity *η* on the hematocrit *φ* according to several selected formulas from Table 1.

Specially for blood, a semi-empirical, more precise formula

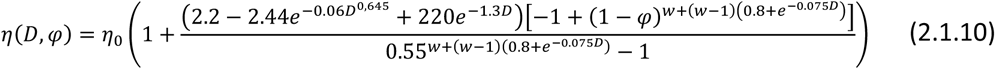

was developed by Pries (36), where *D* represents the tube diameter (in μm) and the factor *w* = 1/(1 + *D*^12^/10^11^). In case of a rapid flow in a small vessel, the erythrocytes, as very large ‘particles’, are concentrated in the middle of the vessel, which decreases the effective blood viscosity (known as Fåhraeus-Lindqvist effect (1, 2)). Therefore, the effective viscosity depends not only on the hematocrit *φ*, but also on the vessel diameter *D*. The Pries formula takes account of this behavior and can be regarded as most reliable. In small vessels, viscosity is less sensitive to hematocrit, which is reflected in the curve for small vessel diameters (50 μm) in Fig. 1, lying below that for large diameters.

The oxygen content in one liter of fully saturated blood, *C*_*ox*_, is proportional to the hematocrit *φ*, with the proportionality constant *k*

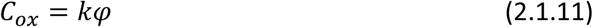

The whole organism oxygen supply, further denoted as J_ox_, can be expressed as

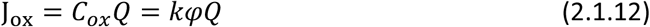

In the following sections, two constrained optimization models of oxygen supply will be described – a model with a constraint of constant driving pressure and a model with a constraint of constant cardiac power.

### 2.2. Optimization of oxygen supply under constant driving pressure

Optimization with constant driving pressure is similar to the optimization performed by our group formerly (1, 2). Only the main ideas of that work will be summarized in this section.

With the substitution for *Q* from Eq. 2.1.2 to Eq. 2.1.12 we obtain J_ox_ as a function of Δ*p* and *φ*

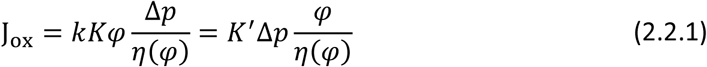

The product *kK* was replaced with a new constant *K*′.

For a given driving pressure Δ*p*, J_ox_ achieves maximum at

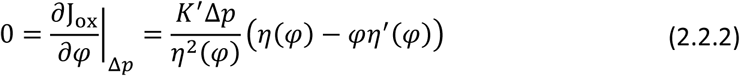

which results in a condition for the optimal hematocrit 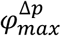

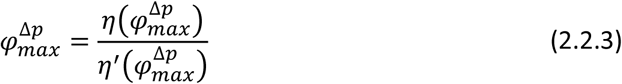

That this leads to a maximum can be confirmed with the negative value of the second derivative.

The different optimal values 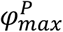 following from various formulas for *η*(*φ*) are given in Table 2 (left column), ranging mostly from 0.3 to 0.4. Selected plots of J_ox_ as a function of *φ* under constant driving pressure are shown in Fig. 2. Fig. 3 illustrates the influence of the vessel diameter, based on the Pries formula. For small vessels, the lower effective viscosity results in a higher optimal hematocrit, reaching the optimal value of more than 0.5 for vessel diameters below 50 μm. For large diameters the Pries formula provides approximately the same optimal value of 0.39 as Saitô’s formula. It is worth noting that the optimization in the model of Farutin et al. (3) was also performed under constant driving pressure and yields a similar optimal hematocrit value of about 0.4.

**TABLE 2.** Comparison of optimal values of the hematocrit, optimized under constant driving pressure (left column) and constant cardiac power (right column)

| Viscosity formula | $\varphi_{max}^{\Delta p}$ | $\varphi_{max}^P$ |
| --- | --- | --- |
| Mooney | 0.234 | 0.344 |
| Quemada | 0.333 | 0.505 |
| Krieger-Dougherty | 0.286 | 0.491 |
| Gillespie | 0.302 | 0.474 |
| Saitô | 0.387 | 0.603 |
| Arrhenius | 0.400 | 0.800 |
| Pries D = 10 $\mu\text{m}$ | 0.608 | 0.731 |
| Pries D = 1000 $\mu\text{m}$ | 0.393 | 0.627 |

**FIGURE 2.**
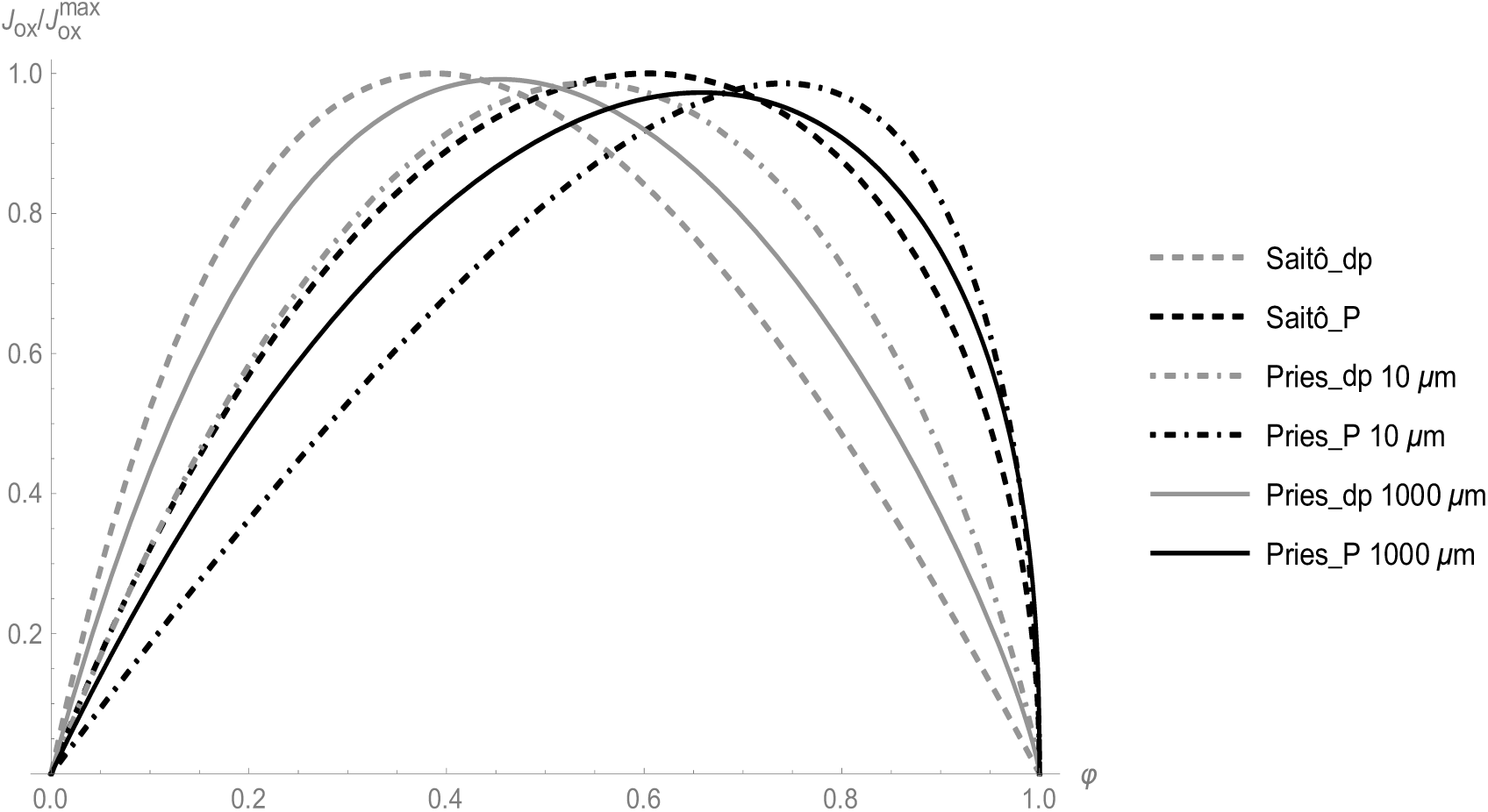
Dependence of J_ox_ on *φ* for several selected formulas for *η*(*φ*). Gray curves - calculation under constant driving pressure (_dp), black curves – calculation under constant cardiac power (_P). J_ox_ is plotted as a fraction of maximal J_ox_ of the respective curve.

**FIGURE 3.**
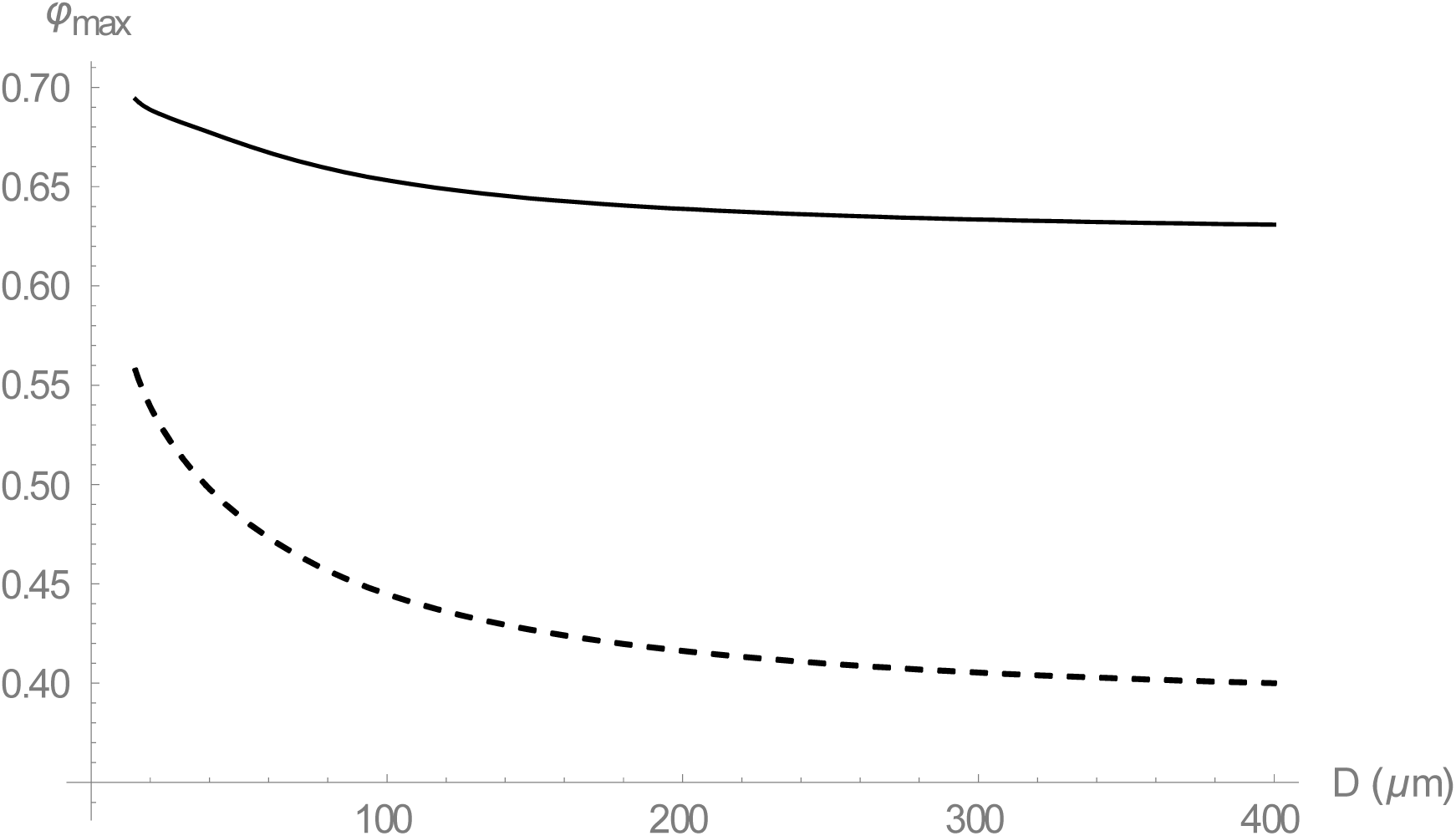
Dependence of the optimal hematocrit *φ*_*max*_ on the vessel diameter D, based on the Pries formula for the optimization under constant driving pressure (dashed curve) and constant cardiac power (full curve).

### 2.3. Optimization of oxygen supply under constant cardiac power

Now we modify the circulation model so that the physical power of the heart *P* rather than the driving pressure Δ*p* remains constant.

The physical power *P* of the heart is defined as the work performed by the heart in a unit time. Thus,

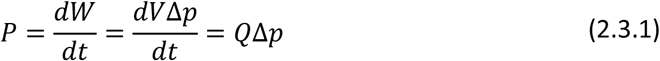

where *W* is the work of the heart and *V* the ejected blood volume.

With a substitution from Eq. 2.1.2 we obtain

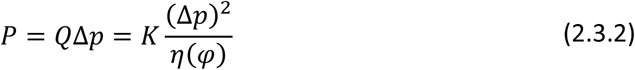

Substituting Δ*p* in Eq. 2.1.2 we obtain *Q* as a function of *P* and *φ*

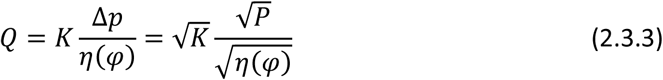

For a given cardiac power *P* the blood flow depends on the inverse square root of viscosity, contrary to just inverse linear dependence in case of constant driving pressure, as follows from Eq. 2.1.2. This makes the blood flow less dependent on viscosity. A lower sensitivity of blood flow to hematocrit at exertion was indeed found in experiment (7).

For J_ox_, it follows

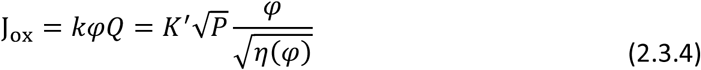

where the constant terms were replaced with a new constant *K*′.

For a given cardiac power *P*, the J_ox_ achieves maximum at

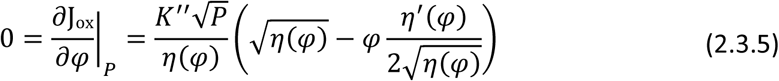

which results in a condition for 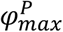

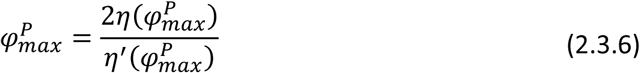

Hence, the 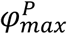 represents the hematocrit that maximizes the oxygen supply under a given cardiac power. Again, this can be confirmed with the negative value of the second derivative.

As in the previous subsection, various formulas for *η*(*φ*) provide different optimal hematocrit values 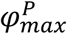, given in Table 2 (right column), ranging mostly from 0.5 to 0.7. Selected plots of J_ox_ as a function of *φ* at a given cardiac power are shown in Fig. 2. Fig. 3 illustrates the influence of vessel diameter, based on the Pries formula. For small vessels the optimal hematocrit increases up to 0.7, for big vessels we obtain an optimal value of around 0.6, just as from Saitô’s formula.

### 2.4. Relationship between 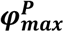 and 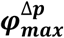

As seen in Table 2, the optimal hematocrit under constant cardiac power is always higher than the optimal hematocrit under constant driving pressure, that is

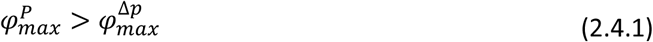

This can be proved in a general way. The implicit Eqs. 2.2.3 and 2.3.6 can be rearranged to

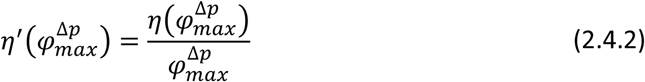

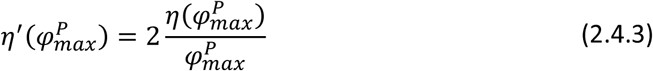

These equations have the following geometric interpretation (Fig. 4). The tangent to the function *η*(*φ*) in the point 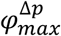 (dash-dotted line) intersects the abscissa in the origin of coordinates. The tangent in the point 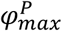 (dashed line) intersects the x-axis in the point 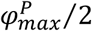. As the function *η*(*φ*) is positive, monotonic increasing and strictly convex, inequality 2.4.1 must always hold.

**FIGURE 4.**
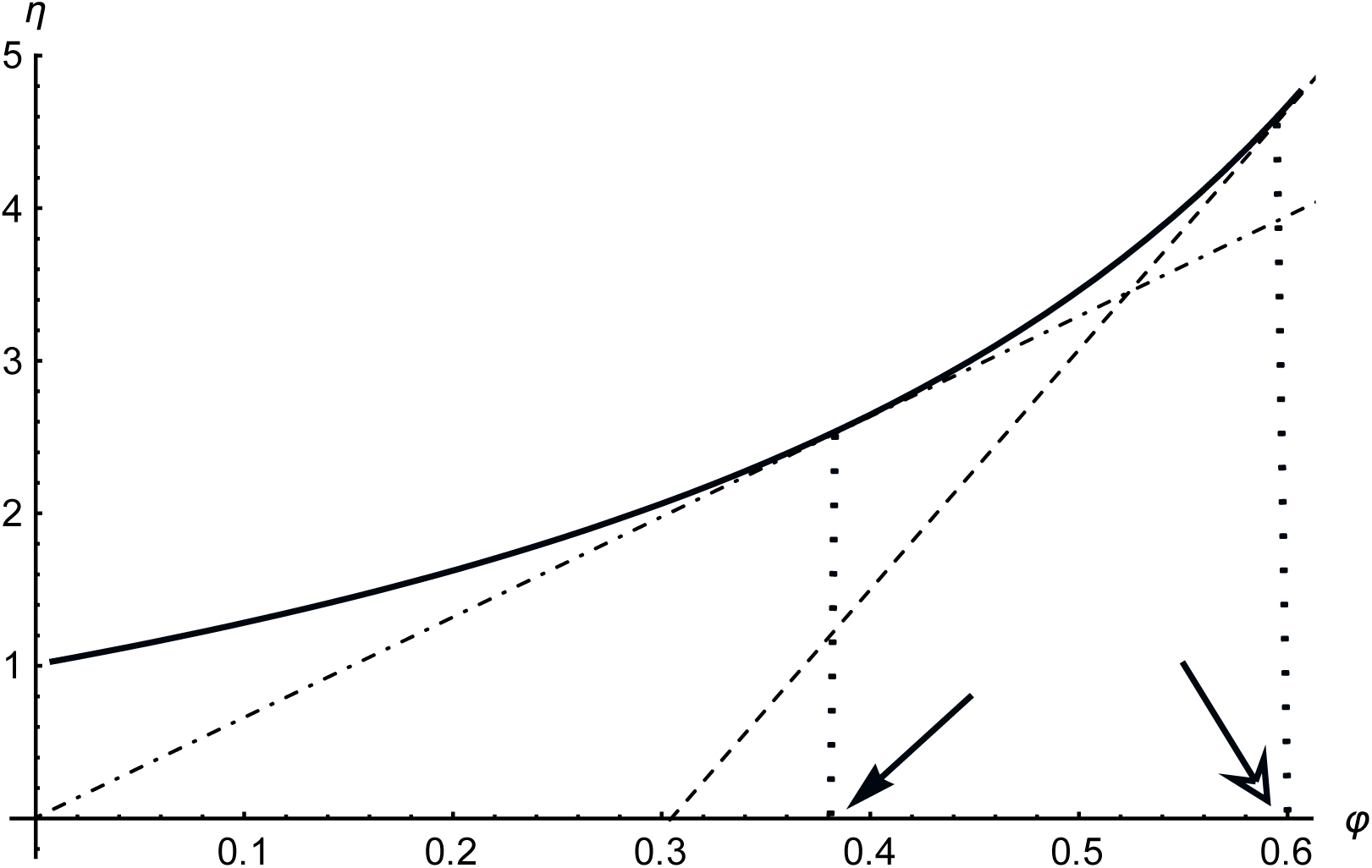
Geometric interpretation of 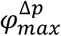 (full arrow) and 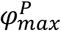 (empty arrow). Full curve –function *η*(*φ*), dash-dotted line - tangent in the point 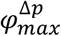, dashed line – tangent in the point 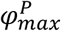. For further explanations, see text.

Intuitively, the higher optimal value can be explained with a weaker dependence of blood flow on the viscosity at given cardiac power – a dependence on the inverse square root of viscosity contrary to the inverse proportionality, as noted above.

## 3. Discussion

We have presented two models of global oxygen supply, which differ in their constraints – notably constant driving pressure and constant cardiac power. We have shown that the oxygen supply in the latter case achieves maximum at higher hematocrit. Particularly, the optimal hematocrit under the constraint of constant driving pressure ranges from 0.3 to 0.5 and under constant cardiac power from 0.5 to 0.7. The increase in viscosity due to higher hematocrit is partly compensated by an increase in the driving pressure.

It is worth mentioning that there exist problems equivalent to the optimization problems under study, which lead to the same optimal hematocrit values but have a more intuitive physical interpretation. The equivalence can easily be shown by rearrangement of the equations or by the method of Lagrange multipliers. First, maximizing oxygen supply under constant driving pressure is equivalent to minimizing the driving pressure under constant oxygen supply. Second, maximizing oxygen supply under constant cardiac power is equivalent to minimizing cardiac power under constant oxygen supply.

We now discuss the physiological relevance of the calculated results and their relation to other studies. The first model with the constant driving pressure yields optimal hematocrit values near the normal values of many animals, such as cat, pig, orangutan, chimpanzee and killer whale (1). Guyton et al. (5) changed, in experiment, the hematocrit in dogs at rest with the use of blood transfusion without any change in blood volume. The blood pressure remained almost unchanged during the experiment. The highest oxygen supply was found at the hematocrit of 0.4. Thus, the first model provides a good explanation of the resting value of the optimal hematocrit.

Different from the situation at rest are the circulatory conditions at an extreme exercise, e.g. during high-performance sport. It was shown that the heart achieves its maximum power in that case and is a limiting factor for the cardiac output (37). Under the plausible assumption that a maximal cardiac power implies that it remains constant over a certain period, the optimization corresponds to our second model. Our model then predicts the optimal hematocrit to range from 0.5 to 0.7, which is in a good agreement with 0.58 in the study of Schuler (21) or 0.6 of hunted horses as an example of animals called natural blood dopers (8).

Thus, the second model could provide the missing theoretical explanation why the optimal hematocrit should be increased at an extreme exertion, although the normal hematocrit is optimal at rest. At an extreme exertion, but not at rest, the energy output of the heart, which equals the energy loss in the circulation, is the limiting factor for the performance. To minimize energy loss, the hematocrit must be higher than normal, as follows from the equivalent problem of minimizing cardiac power under constant oxygen supply mentioned above. Most of the energy is lost on the level of small arterioles with a diameter of 50-100 μm. For this diameter the calculation based on the Pries formula provides the optimal value of 0.65, which is very close to the experimental values of 0.58 or 0.6 mentioned above.

There are, however, several drawbacks of the presented explanation. Firstly, the cardiac power, i.e. the mechanical energy output from the heart, is not necessarily the most relevant constraint at exertion. Another limiting factor, which reaches maximum at exertion, could be the energy (or oxygen) consumption of the heart, which is mainly determined by the coronary perfusion and heart metabolism. If the heart work efficiency, i.e. the percentage of energy consumption converted to mechanical work, was constant, both cardiac energy output and energy consumption would be equivalent constraints in our second model. It was shown, however, that the heart work efficiency markedly declines with rising arterial blood pressure (38) and that cardiac power slightly increases as hematocrit is increased (8). Thus, our second model describes an idealized, extreme case. The true optimal value is likely to be situated between both extreme models. An interesting analogy is a bicycle with gear shift. When using a high gear, the pedals need be moved with low velocity only, yet with high force. There is an upper bound on the force generated by the leg muscles and also by the heart. It is then easier to switch to a lower gear, and so it is to pump the blood at a lower hematocrit. The danger of extreme high blood pressure upon blood doping can be documented by the death of 18 Dutch and Belgian cyclists from 1987-1990 (39).

Secondly, Gaehtgens (6) studied the dependence of oxygen supply on hematocrit in an isolated working skeletal muscle of a dog that was artificially perfused with constant perfusion pressure. He found optimal hematocrit values of 0.5-0.6. The optimal value for a resting skeletal muscle was found to be 0.3-0.4. Contrary to our models, both the resting and exertional optimal values were determined under the constraint of constant perfusion pressure. The authors explain the lower resting optimal hematocrit with a possibility of compensatory vasodilation upon increase in hematocrit at rest (see Fig. 5), but not during exercise, because the arterioles of intensely working muscle are already fully dilated at normal hematocrit. This explanation is, however, questionable because a compensatory vasodilation would make the blood flow-hematocrit curve flatter at rest, at least for hematocrit higher than normal.

**FIGURE 5.**
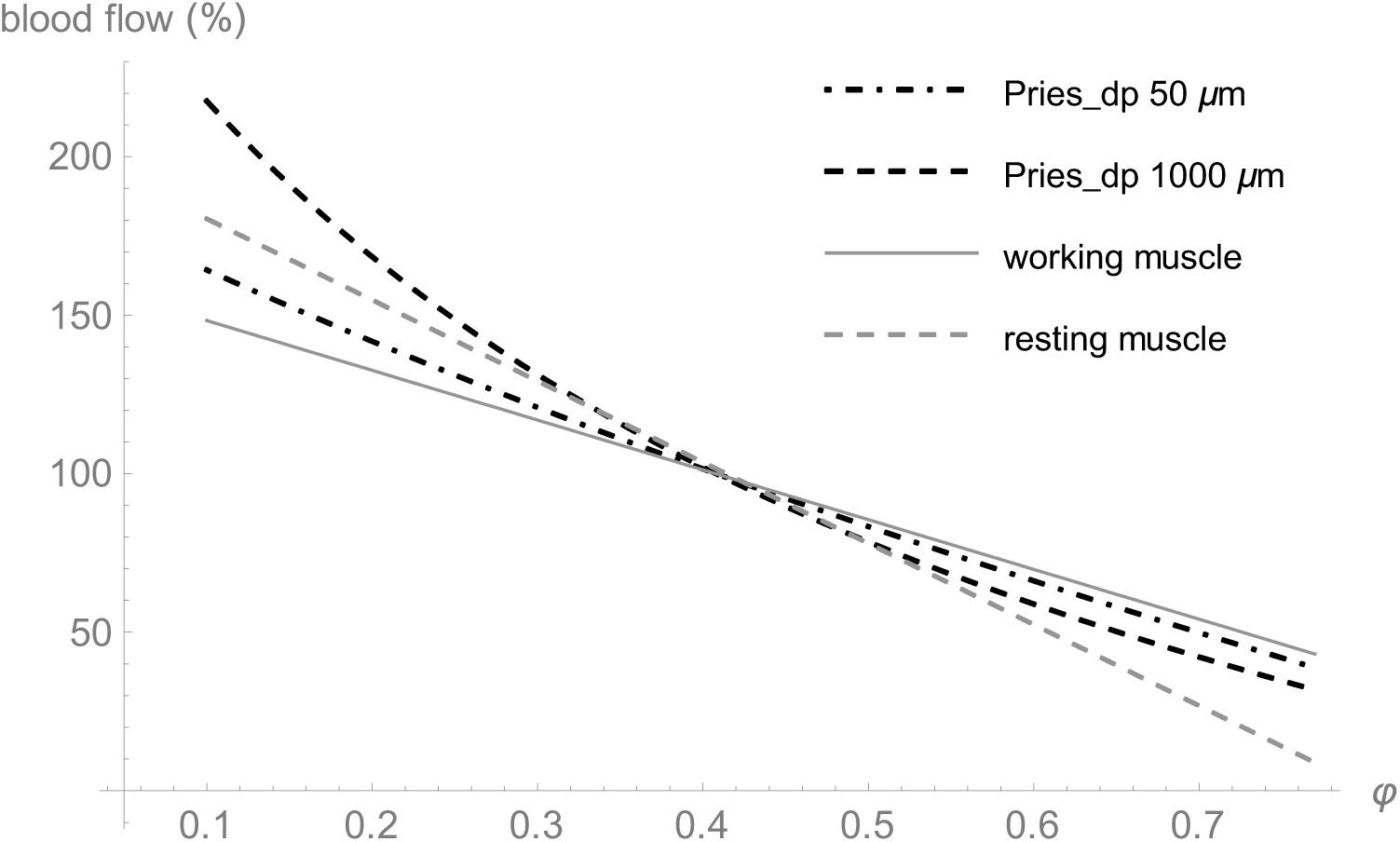
Comparison of experimental data from the study by Gaehtgens (6) with our model predictions under the constraint of constant perfusion pressure – dependence of relative blood flow on hematocrit. Blood flow of 100 % corresponds to a hematocrit of 0.4 for all curves. Dashed black line – model prediction using the Pries formula for a diameter of 1000 μm, dot-dashed black line - model prediction using the Pries formula for a diameter of 50 μm, dashed gray line – experimental data for resting skeletal muscle (fitted in (6) by the linear function *Q = 206 - 256 φ*), full gray line – experimental data for working skeletal muscle (*Q = 164 - 157 φ*).

In Fig. 5 we compare experimental data – the dependence of relative blood flow on hematocrit - taken from Fig. 6 of Gaehtgens’ study (6) with predictions of our first model under the constraint of constant perfusion pressure. For the calculations, we used the Pries formula for small (50 μm) and large (1000 μm) vessels. That formula includes the influence of the vessel diameter on the effective blood viscosity (i.e. the Fåhraeus-Lindqvist effect). As seen from Fig. 5, our predictions for small vessels are in a very good agreement with experimental data for working skeletal muscle in a broad range of hematocrit. In this case, arterioles are supposed to be fully dilated in the whole range of hematocrit, which eliminates any compensatory changes of vascular resistance, just as in our model. Thus, we can explain most of the results of the study of Gaehtgens with our first model, without any consideration of changing vasodilation. Similarly, Fig. 6 shows a very good agreement between the normalized experimental data – the dependence of relative oxygen supply on hematocrit - for the working muscle (black points) taken from Fig. 7 of (6) and our prediction using the Pries formula for small vessel diameter. Both the model and experiment provide the same optimal hematocrit value of 0.5.

**FIGURE 6.**
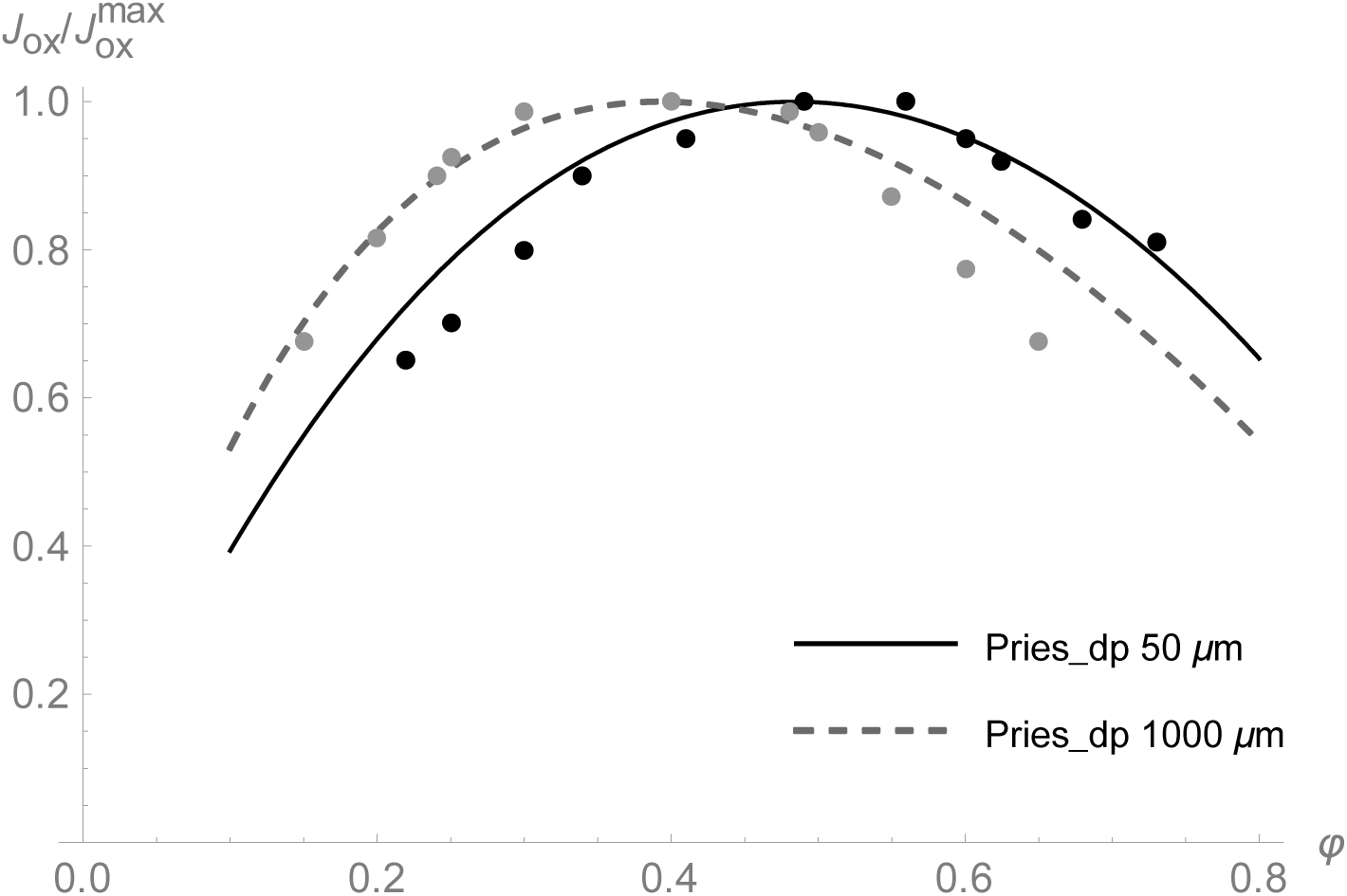
Comparison of normalized experimental data from the studies by Gaehtgens (6) and Guyton (5) with our model predictions under the constraint of constant driving pressure – dependence of oxygen supply on hematocrit. Full black line – model prediction using the Pries formula for diameter of 50 μm, dashed gray line - model prediction using the Pries formula for diameter of 150 μm, black points – experimental data for working skeletal muscle (taken from Fig. 7 of (6)), gray points – experimental data in resting situation (taken from Fig. 3 of (5)).

The agreement of experimental data under resting conditions (gray points in Fig. 6) with our prediction is very good as well (Fig. 5 and Fig. 6). Just the prediction for very low or very high hematocrit values agrees with experimental data less exactly, which could be explained by compensatory vasodilation at extreme values of hematocrit, not considered in our model. The experimental optimal hematocrit value of 0.4 matches with the predicted one accurately (Fig. 6). It is questionable why this agreement applies to a vessel diameter markedly larger than the diameter of arterioles. As shown by Guyton (40), the resistance to venous return rather than arteriolar resistance determines the resting cardiac output, apart from the mean systemic filling pressure. Considering the relatively large diameter of small venules and veins, the coincidence between experimental data and the prediction for big vessel diameters is not surprising.

The agreement of our predictions (based on the Pries formula) for narrow and wide vessels with experimental data at exertion and rest is surprising because arterioles are supposed to be dilated at exertion. We explain this with a mechanical effect of muscle contractions that may compress arterioles and propel blood through the microcirculation and mitigate in this way the influence of hematocrit on viscosity (7). In a similar way, the Fåhraeus-Lindqvist effect mitigates the influence of hematocrit in very small vessels. Note that it is justified to compare experimental data taken from isolated muscle and our prediction for the whole circulation, because of comparable relative changes in cardiac output and blood flow in isolated muscles in dependence on hematocrit (41).

In spite of the very good agreement of the predictions at constant pressure, we maintain that, from a physiological perspective, the constraint of constant perfusion pressure does not correspond well with the heart as a limiting factor at extreme exertion, as described above. Therefore, we consider the second model under constant cardiac power as a valid explanation of the positive effect of the increased hematocrit at exertion, at least as a tendency. The real optimal value is likely to be between both extreme theoretical values.

Experimental data suggest that under normal (resting) conditions, oxygen uptake by the muscles is smaller than the amount delivered by circulation (43). This is not in contradiction to our first optimality criterion because the muscle cannot take up, anyway, 100 % of the oxygen delivered. Moreover, tissues should not take up all the oxygen because of their arrangement in series; part of the oxygen should be left for subsequent cells. Maximizing oxygen delivery per time is in any case favourable, especially in tissues only involving a low density of capillaries, even if other tissues are exposed to an excess of oxygen. Alternatively, we can explain the relevance of the optimality criterion by its equivalent problem of minimizing the pressure difference (which alleviates the effort by the heart) at constant oxygen supply.

It is worth mentioning that even a simple model, which does not consider the many complex effects such as non-Newtonian behavior, deformation and aggregation of erythrocytes, vasodilation, oxygen consumption of the heart etc., can describe the observations fairly well. In future studies, it is of interest, to extend the model by including such effects.

An interesting question is: why did evolution set the hematocrit to the optimal value at rest and not to that at exertion? Probably the long-term risks of the increased hematocrit (e.g. increased blood pressure with a consequent chronic heart failure or increased risk of a thrombosis) may have outweighed the short-term advantages during an extreme exertion. As mentioned above, some animal species, such as dogs and horses, called natural blood dopers, make use of the high hematocrit during exertion, without facing a long-term risk. It is worth noting that both dogs and horses are companions of man used for hunting for a long time and could be selected for blood doping. This is in line with the observation that the top aerobic speed (speed of endurance running) of both animals is about 40 km/h^-1^ and, thus, maximum among mammals (44).

## 4. Conclusions

Our calculations predict optimal hematocrit values of 0.5-0.7 for intense exertion, which is in good agreement with experimental observations for natural blood dopers such as horses (7).

The above calculations can be promising for future applications in personalized medicine. For example, for the therapy of patients with limited cardiac power, as in the cardiogenic shock (45, 46), calculating the optimal hematocrit value could be helpful. The treatment goal for such patients would be to supply a given amount of oxygen, but to minimize necessary cardiac power. A special application may concern the treatment of heart diseases in highlanders of different populations (47).

## Author Contributions

M.S. conception and design of research; M.S., H.S. and S.S. analyzed data; M.S. prepared figures; M.S., H.S. and S.S. drafted manuscript; M.S., H.S. and S.S. edited and revised manuscript; M.S., H.S. and S.S. approved final version of manuscript

## Acknowledgements

We thank Bashar Ibrahim (Jena) for helpful discussions and three anonymous referees for very helpful suggestions.

## Notes

### Competing Interest Statement

The authors have declared no competing interest.

